# Transcription activity contributes to the firing of non-constitutive origins in *Trypanosoma brucei*

**DOI:** 10.1101/398016

**Authors:** Marcelo S. da Silva, Gustavo R. Cayres-Silva, Marcela O. Vitarelli, Paula A. Marin, Priscila M. Hiraiwa, Christiane B. Araújo, Andrea R. Ávila, Marcelo S. Reis, Maria Carolina Elias

## Abstract

The cosynthesis of DNA and RNA potentially generates conflicts between replication and transcription, which can lead to genomic instability. In trypanosomatids, eukaryotic parasites that perform polycistronic transcription, this phenomenon and its consequences have not yet been investigated. Here, using equations and computational analysis we demonstrated that the number of constitutive origins mapped in the *Trypanosoma brucei* genome is close to the minimum required to complete replication within S phase duration. However, taking into account the location of these origins in the genome, the replication in due time becomes virtually impossible, making it necessary to activate non-constitutive origins. Moreover, computational and biological assays pointed to transcription being responsible for activating non-constitutive origins. Together, our results suggest that transcription action through conflicts with replication contributes to the firing of non-constitutive origins, maintaining the robustness of S phase duration. The usage of this entire pool of origins seems to be of paramount importance for the survival of this parasite that infects million people around the world since it contributes to the maintenance of the replication of its DNA.

## INTRODUCTION

DNA replication is an essential biological process because it is the basis of genetic inheritance. In cellular organisms, the earliest step in DNA replication is the establishment of origins, which are defined as genomic loci where DNA synthesis initiates. In eukaryotes, replication initiation is preceded by the binding of an initiator called the Origin Recognition Complex (ORC) (1). This initiator establishes replication initiation through the recruitment and activation of the replisome; hereafter, we refer to this phenomenon as “origin firing”. An origin firing yields two replication forks, which proceed in opposite directions (bi-directional movement) following a DNA replication rate that varies according to the cell type (1–3). The time required to complete DNA replication on all chromosomes determines the S phase duration, which can be one of the primary methods to regulate cell cycle progression in eukaryotes (4, 5). Regarding the number of origins per chromosome, bacteria typically have a single origin, whereas eukaryotes and archaea generally have multiple origins per chromosome (1, 6), although the exact number varies according to cell type and the cellular environment.

During the G1 phase of the cell cycle, all potential replication origins are licensed, *i.e*., the loading of DNA helicase (namely, MCM_2-7_) is carried out (6, 7). However, during the progression of a single cell cycle, only a subset of these origins activates DNA synthesis. The choice of origins that will be activated in the cell cycle varies from cell to cell, which implies a flexibility regarding origin usage (6). According to their different usages, replication origins can be classified into three categories: constitutive, which are always activated in all cells of a given population; flexible, whose usage varies from cell to cell; and dormant, which are not fired during a normal cell cycle but are activated in the presence of replication stress (6).

An important phenomenon that may contribute to replication stress is replication-transcription conflicts. These clashes are a powerful source of genomic instability that can generate DNA double-stranded breaks (DSBs), which in turn impair DNA synthesis (8–10). However, whether replication-transcription conflicts contribute to the activation of origins to avoid lengthening S phase and consequent replication impairment is a question that remains open.

In trypanosomatids, unlike eukaryotic model organisms, the majority of genes are organized into large polycistronic gene clusters (11), which makes us wonder whether these organisms regulate transcription and/or replication in order to mitigate potential collisions during S phase. Trypanosomatids encompass human pathogens of great medical importance, such as *Trypanosoma* spp. and *Leishmania* spp., which are the causative agents of devastating diseases that threaten millions of people around the world (12, 13). *Trypanosoma brucei*, one of the parasites known as African trypanosomes, has the best-studied DNA and RNA synthesis, although this knowledge is limited relative to that of model eukaryotes (14). The duration of the by using a most sensitive thymidine analog, 5-ethynyl-2’-deoxyuridine (EdU), to monitor DNA replication, although there are still no similar assays for *T. brucei* TREU927 (15). The number of DNA replication origins per chromosome and the replication rate are a matter of debate according to the technique used to obtain these data and the choice of either *T. brucei* Lister strain 427 or TREU927 (3, 14, 16, 17). Even with its peculiar feature of performing polycistronic transcription in large gene clusters, thus far there have been no studies of replication-transcription conflicts in trypanosomatids.

In this work, we investigated the dynamics of origins usage during S phase and determined whether the action of the transcription machinery contributes to the activation of non-constitutive origins in *T. brucei* strain TREU927. To this end, we used EdU to monitor DNA replication and more accurately estimate S phase duration. Then, based on the chromosome size, specific replication rates, and the precise S phase duration, we developed a mathematical formula to estimate the minimum number of DNA replication origins required to duplicate an entire chromosome within S phase. After applying this formula to each chromosome, we compared the minimum number of origins (*mo*) obtained with the number of constitutive origins mapped by the MFA-seq technique, and we found that these values were very similar. However, when we used another formula that takes into account replication rates and the positions of mapped constitutive origins, we determined that these mapped origins were not sufficient to allow complete replication within the S phase duration, largely due to their localization in each chromosome. We then designed and simulated stochastic computational dynamic models involving origin firing in the presence or absence of polycistronic transcription gene clusters during S phase. These simulations suggested the robustness of S phase duration relative to increasing levels of transcriptional activity, which were compensated by increased replication initiation. Based on these results, we developed a hypothesis that could satisfactorily explain the observed phenomenon: if there is transcription during S phase, then the cosynthesis of DNA and RNA can generate collisions between the two associated types of machinery, activating additional origins (flexible and/or dormant origins) and consequently increasing the pool of origins used to complete S phase. Using a run-on assay, we showed the transcription landscape of the *T. brucei* TREU927 cell cycle, verifying that there is transcription during S phase. Moreover, we verified the presence of γH2A foci (a DNA damage biomarker) and R-loops during S phase, which decreased after transcription inhibition, suggesting a role for transcription action in the formation of DNA lesions and DNA:RNA hybrids. In addition, we showed that γH2A levels colocalize with R-loops in late S/G2 phase and decrease after transcription inhibition. Finally, using a DNA combing technique, we detected fewer numbers of activated origins after transcription inhibition. Additionally, we measured the length of S phase and observed that it remained unchanged.

Taken together, our results indicate the firing of non-constitutive origins as a result of transcription activity, possibly through replication-transcription conflicts. In other words, our results suggest that the action of transcription machinery contributes to the activation of origins above the theoretical minimum to maintain the robustness of S phase duration in *T. brucei*. This work is the first to show evidence of replication-transcription conflicts in trypanosomatids and their consequences for genome maintenance in these human pathogens of great medical importance.

## MATERIALS AND METHODS

### Trypanosomatid culture and growth curves

*T. brucei* procyclic forms (TREU927) were cultured at 28 °C in SDM79 medium supplemented with 10% (v/v) fetal bovine serum. To generate growth curves, parasite culture was initiated with 1 × 10^6^ cells.mL^-1^ (daily curves) or with 10 × 10^6^ cells.mL^-1^ (hourly curves), and cells were harvested and counted until they reached the stationary phase.

### EdU incorporation assays and ‘click’ chemistry reaction

Exponentially growing parasites were incubated with 100 µM 5-ethynyl-2’-deoxyuridine (EdU) (ThermoFisher Scientific) for the time required according to each assay (ranging between 1–2.25 h) at a species-specific temperature (28 °C for *T. brucei*). The parasites were then harvested by centrifugation at 2,500 *g* for 5 min, washed three times in 1x PBS (137 mM NaCl, 2.7 mM KCl, 10 mM Na_2_HPO_4_, and 2 mM KH_2_PO_4_, pH 7.4), and the pellet was resuspended in 200 µL of the same buffer solution. Afterward, 100 µL of the cell suspension was loaded onto poly-L-lysine pretreated microscope slides (Tekdon), fixed for 20 min using 4% sterile paraformaldehyde (Merck) diluted in 1x PBS, washed three times with 1x PBS, and then washed three times with 3% BSA (Sigma-Aldrich) diluted in 1x PBS. Then, parasites were permeabilized for 10 min with 0.1% sterile Triton X-100 (Sigma Aldrich) diluted in 1x PBS, washed three times with 1x PBS, and then washed three times with 3% BSA in 1x PBS. To detect incorporated EdU, we used Click-iT EdU detection solution for 45 min and protected the reaction from light. The Click-iT EdU detection mix solution consisted of 25 µL of 500 mM ascorbic acid (C_6_H_8_O_6_), 5 µL of 100 mM copper sulfate (CuSO_4_), 2.5 µL of Alexa fluor azide 488 (ThermoFisher Scientific), and 467.5 µL of distilled water (for details about the EdU procedure, see Salic and Mitchison, 2008) (18). Finally, the parasites were washed five times with 1x PBS. Vectashield Mounting Medium (Vector) containing 4’,6-diamidino-2-phenylindole dihydrochloride (DAPI) was used as an antifade mounting solution and to stain nuclear and kinetoplast DNA. Images were acquired using an Olympus Bx51 fluorescence microscope (100x oil objective) attached to an EXFO Xcite series 120Q lamp and a digital Olympus XM10 camera with camera controller software Olympus Cell F (Olympus, Japan). Images were further analyzed using ImageJ software (National Institutes of Health) to count the numbers of EdU-positive parasites, and the percentage of proliferating parasites was calculated for each sample relative to the total number of DAPI-positive parasites.

### Analysis of the cell cycle

Formaldehyde-fixed and DAPI-stained exponentially growing procyclic forms of *T. brucei* TREU927 were examined under an Olympus BX51 fluorescence microscope (Olympus) (100x oil objective) to observe the profile of organelles containing DNA (nucleus and kinetoplast). To estimate the duration of cytokinesis (C), we used the Williams (1971) Equation (19):

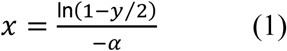

where x is the cumulative time within the cycle until the end of the stage in question, y is the cumulative proportion of cells up to and including the stage in question (expressed as a fraction of one unit), and α is the specific growth rate.

To estimate the G2+M phases, we added EdU in each culture medium containing the parasites for analysis and collected them every 15 min until a parasite containing two EdU-labeled nuclei (2N2K) was observed.

The duration of S phase was estimated according to the Stanners and Till (1960) Equation (20):

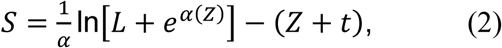

where *L* is the proportion of cells exhibiting EdU-labeled nuclei, α = ln 2.T^-1^ (*T* = doubling time expressed in hours), Z = G2 + M + cytokinesis, and *t* is the duration of the EdU labeling period in hours. Finally, the duration of the G1 phase was estimated by the difference between the doubling time and the sum of the remaining phases.

### Estimation of the minimum number of origins required to complete S phase

To determine the minimum number of origins needed to replicate an entire chromosome, we developed a mathematical formula. This formula uses the S phase duration (*S*), the size of the chromosome in question (*N*), and the replication rate (*v*). The lower bound *mo* for the number of origins required to replicate an entire chromosome is given by:

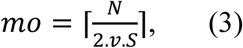

Of note, if the right-hand side of the formula results in a fraction of a unit, then the next higher integer unit must be taken as the result of the formula, which is represented by the ceiling function (⌈ ⌉). In Theorem S1 (Supplementary Material), we prove the correctness of this equation.

We used up-to-date data available in TriTrypDB database (www.tritrypdb.org) and data reported in recent studies as parameters for the formula (3, 17). We then compared the obtained results with the origins mapped by the MFA-seq technique, presented in another study (16).

### AT content enrichment analysis and probability landscape for origin firing

AT-content distribution (Supplementary Figure 1, red lines) was computed for *T. brucei* TREU927 through an in-house Perl program that split each chromosome into sets of 1,000 bp-sized bins and computed the AT content of each bin.

**Figure 1.**
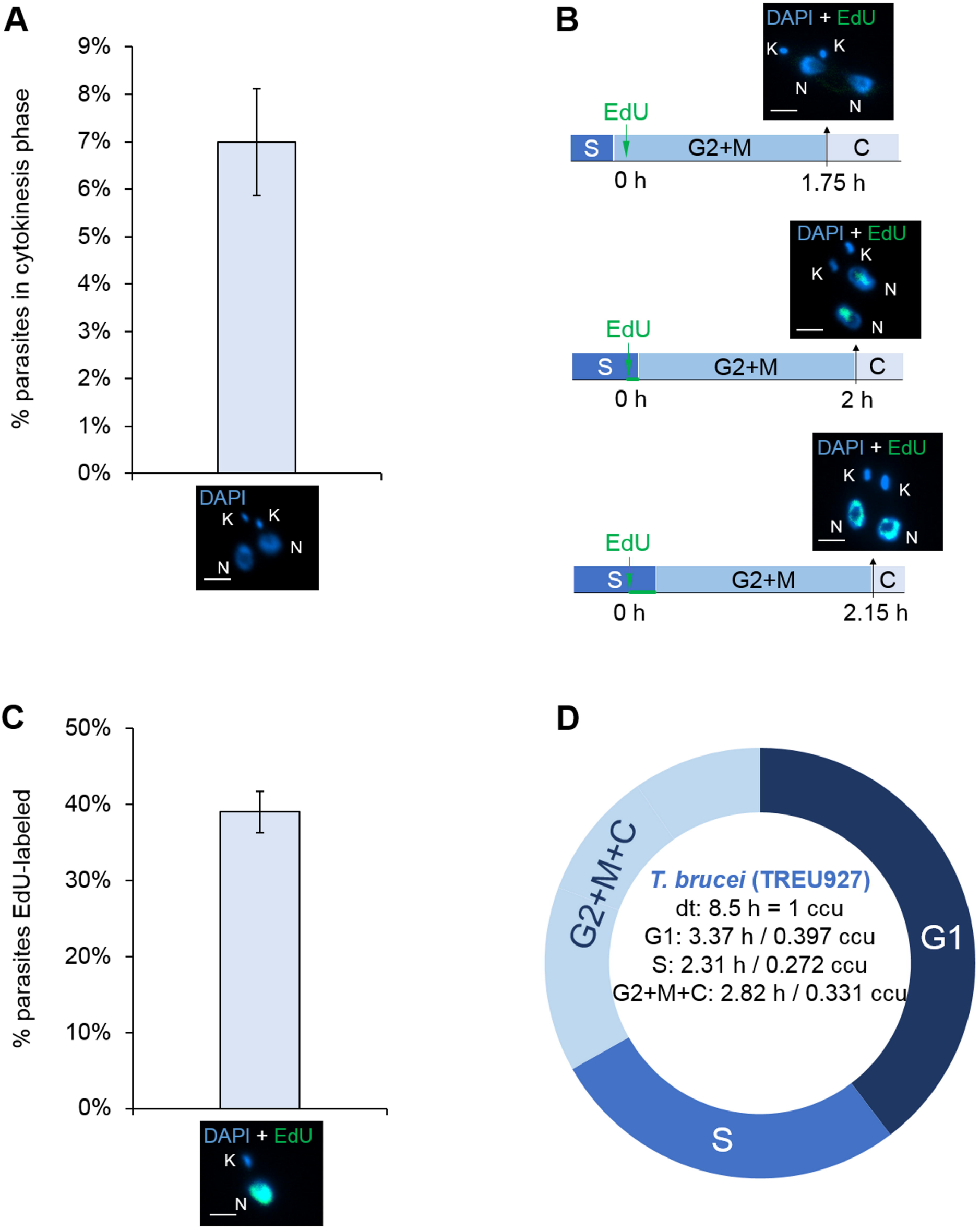
Estimation of S phase duration in *T. brucei* TREU927. **A**. DAPI-labeled parasites (2K2N) were used to measure the percentage of parasites in cytokinesis, which was estimated to be 6.99 ± 1.13%. Error bars represent SD. The scale bars for the fluorescence images correspond to 2 µm. This value was used in equation 1 (19) to estimate the cytokinesis phase duration. **B.** To estimate the duration of the G2+M phases, EdU was added to the culture, and parasites were collected every 15 min until parasites containing two EdU-labeled nuclei were observed. This pattern was observed after 2 h. This assay was carried out in triplicate, and in all replicates, we found a parasite containing two EdU-labeled nuclei at the same time. Scale bar = 2 µm. **C**. EdU-labeled parasites 1 h after EdU pulse were used to estimate the percentage of parasites able to uptake this thymidine analog (39 ± 2.7%). Error bars represent SD. Scale bar = 2 µm. This value was used in equation 2 (20) to estimate the S phase duration. These assays were carried out in biological triplicate (n = 358 parasites). **D.** New precise lengths for G1 and S phase duration. ccu means cell cycle unit, where one unit corresponds to the specific doubling time (dt) for each strain.

The probability landscape for origin firing (Supplementary Figure S1, blue lines) was generated by assigning to each base pair a probability of origin firing. This probability was derived from a marker frequency analysis coupled with deep sequencing (MFA-seq) (16), whose processed results were stored in TriTrypDB. For each chromosome, its corresponding MFA-seq data are composed of a thousand equally sized bins. Each bin has a positive real value. Thus, we mapped these real values into a probability value within the interval [0,1]. In doing so, for each chromosome, we carried out a linear transformation in which we took into account the minimum and maximum bin values, thus yielding our probability landscape.

### Analytical solution for deterministic DNA replication with constitutive origins

Initially, we considered only the origins mapped by the aforementioned MFA-seq assay (16), aiming to reconcile those data with the experimentally observed S phase duration (Figure 1) and two replication rates measured previously (3, 17). To this end, we derived an analytical solution for the lower-bound required time for DNA replication that relies solely on a set Θ = {θ_1_, …, θ_|Θ|_} of constitutive origin locations, the chromosome size *N* and the replication rate *v*:

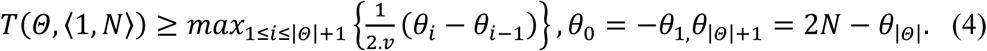

In Theorem S2 (Supplementary Material), we prove the correctness of Equation 4.

### Simulation of a stochastic dynamic model for replication-transcription conflicts

To investigate how putative conflicts between replication and transcription machineries might impact key DNA replication parameters such as S phase duration robustness and mean inter-origin distance (IOD), we designed and simulated a stochastic dynamic model. This model is composed of a set of binary vectors, one vector per chromosome (Supplementary Figure S2). The dimension of each vector was equal to the number of base pairs in that chromosome; therefore, each dimension consisted of a binary flag that assigns whether its corresponding base pair was replicated or not.

In this model, we assume the presence of flexible origins and/or dormant origins with a passive mechanism (*i.e*., origins that fire stochastically). To this end, we modeled the replication process for each chromosome as a Markov chain whose state transition was defined as a function of two parameters: the probability of an origin firing when it is selected at base i in chromosome j (p(i,j)), and the maximum number of replisomes activated simultaneously (*F*), a constraint that was also used in a similar model (21). p(i,j) was estimated through the linear transformation of MFA-seq data (16); this transformation is depicted by the blue lines in Supplementary Figure S1. We also included in the model constitutive transcription, *i.e*., transcription whose levels (i.e., the initiation frequency) remain fairly constant despite the cellular condition. The average transcription rate of RNA polymerase (RNAP) was set to be the same as the replisome replication rate (*v*). Transcription was modeled with periodic firing of RNAP at the beginning of each polycistronic region (Supplementary Figure S3). DNA replication progress was accomplished by updating the binary vectors that represent the replication state of all chromosomes. When a given replication origin is fired, it spawns two replication forks that move in opposite directions (Supplementary Figure S4). The collisions between DNA replication and transcription machineries arise when we simulate together previously described replication and transcription dynamics in the same model (Supplementary Figure S5). In our simulations, we focused our attention on *head-to-head* collisions, in which the replisome and RNAP come from opposite directions. Of note, once we assume that the transcription velocity is equal to replication rate, there are no *head-to-tail* collisions, *i.e*., there is no possibility of an RNAP colliding behind a replisome or vice-versa. The outcome of *head-to-head* collisions was designed following the scenario modeled by Lin and Pasero (2012) (22): if one such event occurs, then there is a fork collapse, and the replisome is released from the chromosome. We carried out several sets of Monte Carlo simulations (30 simulations per set). Each set contained simulations with the same frequency of constitutive transcription (including frequency = 0, that is, no transcription at all) and the same value for parameter F. The obtained results were then averaged (Supplementary Table S2).

Model implementation: To carry out simulations of S phase dynamics in *T. brucei* TREU927, we developed a simulator whose main procedures are presented in Supplementary Figure S6. For each simulation, we constrained the number of available replisomes during a given simulation (parameter *F*) in a manner similar to the procedure adopted in another meticulous study (21).

A simulation starts with all binary vectors (Supplementary Figure S2) filled with zeros (*i.e*., all nucleotides are nonreplicated). After that, at the beginning of each iteration, the simulator verifies whether there are replisomes attached to the chromosomes: if there are, then the simulator advances them, filling the section of the binary vector that was replicated. The simulator also evaluates and solves the replication-transcription conflicts that eventually arise. In the sequence, if the number of activated origins is smaller than or equal to *F -* 2, then one nucleotide of the whole genome is drawn with uniform probability. Origin firing is decided by verifying a given nucleotide probability in the origin firing probability landscape (Supplementary Figure S1). In simulations with transcription, in some iterations, RNAP binds onto the beginning of all polycistronic regions, which depends on a parameter called frequency of constitutive transcription. A simulation ends when the genome is completely replicated or if it reaches a given replication time threshold; the latter might be necessary for replication-transcription conflicts assays since in some simulations the required time for replication completion might be far higher than the mean S phase duration, which means that the simulated organism is not viable.

A priori information was organized using the SQLite relational database manager (www.sqlite.org). The model simulator was implemented in Python 3 programming language (www.python.org) using the NumPy scientific computing library (www.numpy.org). The source code is under the GNU GLPv3 license and can be obtained for free at github.com/LECC-IBU/ReDyMo.

### Nuclear run-on assay to detect transcription throughout the cell cycle

This assay was carried out as described by Hiraiwa et al. (2018) (23), whose protocol is optimized for RNAP II activity, which mainly transcribes messenger RNA (mRNA). Briefly, the parasites were washed with buffer A (150 mM sucrose, 20 mM L-glutamic acid potassium, 20 mM HEPES-KOH pH 7.7, 3 mM MgCl_2,_ 1 mM DTT) and permeabilized with 250 µg/mL LPC (L-α- lysophosphatidylcholine) in the same buffer. Then, the parasites were resuspended in transcription buffer (150 mM sucrose, 20 mM L-glutamic acid potassium, 20 mM HEPES-KOH pH 7.7, 3 mM MgCl_2,_ 1 mM DTT, 0.6 mg/mL creatine kinase, 25 mM creatine phosphate, 8U RNase Out, 10 µg/mL leupeptin, 4 mM ATP, 2 mM GTP, 2 mM CTP, 2 mM BrUTP), and *in vivo* transcription was performed at 28 °C for 15 min under gentle agitation. The reaction was stopped by the addition of buffer A, and the parasites were recovered for immunolabeling with anti-bromodeoxyuridine (anti-BrdU antibody) (1:200) and an anti-rat secondary antibody conjugated to Alexa Fluor 488 (1:500). After that, the parasites were resuspended in 1x PBS containing 2 µg.mL^-1^ DAPI and 10 µg.mL^-1^ RNase A for flow cytometry analysis. Data were collected by flow cytometry (FACSCanto II, BD Biosciences, San Jose, CA, USA). The Alexa Fluor 488 fluorochrome (detection of nascent RNA) was excited with a blue laser (488 nm), and the emitted light was collected with a 530/30 bandpass filter. The DAPI stain (DNA content analysis) was excited with a violet laser (405 nm) and collected with a 450/40 bandpass filter. No compensation was needed. The data were analyzed using FlowJo software (BD, USA) version 10.0.1.

Alternatively, an *in vitro* transcription reaction was carried out with parasites previously treated with 75 μg/mL α-amanitin (an irreversible transcription inhibitor) for 4 min to specifically inhibit RNAP II transcription (23).

### Indirect Immunofluorescence (IIF) assays

Exponentially growing procyclic forms of *T. brucei* were harvested by centrifugation (∼ 5.10^6^ parasites) at 2,500 *g* for 5 min, washed three times with 1x PBS, fixed for 10 min using sterile 4% paraformaldehyde (Merck, Germany) diluted in 1x PBS, and washed again with 1x PBS. Of note, for the IIF assays involving γH2A and R-loops, the procyclic forms of *T. brucei* TREU927 were previously treated with α-amanitin and/or treated with ionizing radiation (50 Gy). After that, these samples were treated in the same manner as described previously, *i.e*., harvested by centrifugation, washed, fixed and washed again.

Washed parasites were homogenized in 1x PBS and added for 15 min to Teflon-coated microscope slides (Tekdon). Parasites were then washed three times on slides for two min each with blocking solution (4% BSA in 1x PBS) and permeabilized for 10 min with 0.1% Triton X- 100 diluted in 1x PBS. Samples were then incubated at room temperature for 2 h with antisera solution containing anti-γH2A polyclonal rabbit antibodies (generously provided by Dr. Richard McCulloch, Wellcome Centre for Molecular Parasitology, University of Glasgow) diluted at 1/1000 in 4% BSA, together with anti-R-loop antibody (DNA:RNA hybrid monoclonal mouse antibody, clone S9.6 – Kerafast) diluted at 1/50 in 4% BSA. After that, parasites were washed three times on slides and incubated for 1 h with a goat anti-rabbit IgG secondary antibody conjugated to Alexa Fluor 555 (Life Technologies) diluted at 1/500 in 1% BSA, together with a goat anti-mouse IgG secondary antibody conjugated to Alexa Fluor 488 (Life Technologies), also diluted at 1/500 in 1% BSA. Afterward, parasites were washed five times, and Vectashield Mounting Medium (Vector) containing 4’,6-diamidino-2-phenylindole dihydrochloride (DAPI) was used as an antifade mounting solution and to stain nuclear and kinetoplast DNA. Simple immunofluorescence images were acquired using an Olympus Bx51 fluorescence microscope (100x oil objective) attached to an EXFO Xcite series 120Q lamp and a digital Olympus XM10 camera with Olympus Cell F camera controller software. The intensity of images captured was estimated using Olympus Cell F tools for 100 cells per sample. When necessary, images were superimposed using Olympus Cell F software. Alternatively, double-immunofluorescence images were analyzed using Olympus IX81 microscope and acquired through a Z-series at a thickness of 0.20 μm (ten layers) using a 100X 1.35NA lens, a disk scanning unit (DSU) that provides confocal-like images using a white light, an arc excitation source, and a CCD camera (Olympus). After acquisition, the images were improved by deconvolution using Autoquant software (Media Cybernetics, Rockville, MD, USA), version X2.1.

### DNA combing

Exponentially growing procyclic forms of *T. brucei* (α-amanitin treated and non-treated) were grown to a concentration of ∼ 6.10^6^ parasites.mL^-1^ and sequentially labeled with two thymidine analogs: 5-iodo-2’-deoxyuridine (IdU, Sigma) at 100 μM for 20 min and 5-chloro-2’- deoxyuridine (CldU, Sigma) at 100 μM for 20 min, without an intermediate wash. After labeling, the cells were immediately harvested by centrifugation at 2,500 *g* for 5 min at 4 °C, washed once with cold 1x PBS and resuspended in 100 μL of 1x PBS with 1% low-melting agarose in order to embed cells in agarose plugs. Then, the plugs were incubated in 500 μL of lysis solution (0.5 M of EDTA, pH 8.0, 1% N-lauryl-sarcosyl and 4 μg.mL^-1^ proteinase K) at 50 °C for 24 h. Next, fresh lysis solution was added, and the plugs were incubated for another 24 h. The plugs were carefully washed several times using 0.5 M EDTA, pH 8.0, to propitiate the complete removal of digested proteins and other degradation products. Protein-free DNA plugs were then stored in 0.5 M EDTA, pH 8.0, at 4 °C or used immediately. Plug samples were washed in TE buffer (10 mM Tris-HCl, pH 8.0; 1 mM EDTA pH 8.0), resuspended in 1 mL of 0.5 M MES buffer (2-(N-morpholino) ethanesulfonic acid, pH 5.5) and melted at 68 °C for 20 min. The solution was maintained at 42 °C for 10 min and treated overnight with 2 U of β-agarose (New England Biolabs). After digestion, 1 mL of 0.5 M MES was added carefully to the DNA solution, and then DNA fibers were regularly stretched (2 kb.μm^-1^) on appropriate coverslips as described previously (24).

Combed DNA was fixed onto coverslips at 65 °C for at least 2 h, denatured in 1 M NaOH for 20 min and washed several times in 1x PBS. After denaturing, coverslips containing the DNA fibers were blocked with 1% BSA diluted in 1x PBS. Immunodetection was performed using primary antibodies (mouse anti-BrdU antibody clone B44 – 1/20 dilution, Becton Dickinson; and rat anti-BrdU antibody clone BU1/75 (ICR1) – 1/20 dilution, Abcam) diluted in 1% BSA and incubated at 37 °C in a humid chamber for 1 h. Of note, mouse anti-BrdU reacts with IdU and BrdU (25, 26), and the rat anti-BrdU antibody cross-reacts with CldU but does not cross-react with thymidine or IdU (26). Next, the coverslips were incubated with goat anti-rat Alexa 488 secondary antibodies (1/20 dilution, Life Technologies) and with goat anti-mouse Alexa 568 antibodies (1/20 dilution, Life technologies). Each incubation step with antibodies was followed by extensive washing with 1x PBS. Then, DNA immunodetection was performed using an anti-ssDNA antibody (1/50 dilution, Chemicon) and goat anti-mouse Alexa 350 (1/10 dilution, Life Technologies). Coverslips were then mounted with 20 μl of Prolong Gold Antifade Mountant (ThermoFisher), dried at room temperature for at least 2 h and processed for image acquisition using an Olympus BX51 fluorescence microscope (100x oil objective) attached to an EXFO Xcite series 120Q lamp and a digital Olympus XM10 camera with Olympus camera controller software. When necessary, images were superimposed using the software Olympus Cell F (Olympus, Tokyo, Japan). The observation of longer DNA fibers required the capture of adjacent fields or the use of a fluorescence microscope equipped with a motorized stage that enabled the scanning of slides with high precision. Fibers < 100 kb were excluded from the analysis. The percentage of origins activated during the thymidine pulses was measured manually using ImageJ software. Statistical analyses of origin density were performed using Prism 5.0 (GraphPad). At least two independent combing experiments were performed for the analysis presented in this study.

### S phase length analysis using the measurement of cytokinesis-labeled nuclei

To estimate the duration of S phase, we used an EdU pulse (100 μM during 15 min) in an exponentially growing culture of *T. brucei* (control and α-amanitin-treated). The parasites were then harvested every 15 min, washed three times in 1x PBS, and fixed for 10 min with 3.7% sterile paraformaldehyde (Merck) diluted in 1x PBS. This same approach was also performed in the presence of α-amanitin, *i.e*., in the absence of transcription. Afterward, the parasites were washed, added to a slide (Tekdon) containing 0.1% poly-L-lysine, permeabilized for 10 min with 0.1% sterile Triton X-100 (Sigma) diluted in 1x PBS, and treated using ‘click’ chemistry to detect EdU (for more details, see EdU incorporation assays and ‘click’ chemistry reaction section described previously). After the addition of Vectashield Mounting Medium containing DAPI (Vector) to stain nuclei (N) and kinetoplasts (K), we sealed the slides, and we measured the percentage of parasites containing two labeled nuclei (cytokinesis). The duration of S phase starts when the first cytokinesis-labeled cell appears, extends throughout the period of detection for all cytokinesis- labeled cells and ends when some unlabeled cells appear again. Of note, this assay was carried out according to a protocol described in a previous study (15).

## RESULTS

### EdU allows a more accurate estimation of S phase duration in *T. brucei* TREU92

To investigate the origin usage dynamics under standard situations in *T. brucei* TREU927, we first needed accurate values for S phase duration, which could be obtained from other studies. However, our group recently published a study highlighting significant differences between the thymidine analogs BrdU and EdU, commonly used to monitor DNA replication in most organisms (15). In summary, this study shows that EdU is much more sensitive for monitoring DNA replication than BrdU, and its usage provides a more accurate estimate of the duration of the cell cycle phases G1, S, and G2 (15). Consequently, this study pointed to = skepticism regarding the accuracy of analyses performed to monitor DNA replication using BrdU (with a DNA denaturation step carried out with 2 M HCl) in trypanosomatids. Therefore, to ensure greater accuracy of S phase duration in *T. brucei* TREU927, these analyses had to be redone using EdU.

First, we performed growth curves to estimate the doubling time (Supplementary Figure S7), which was used in Equations 1 and 2 (see materials and methods) (19, 20). In addition to the doubling time, we also estimated the percentage of parasites performing cytokinesis (C), which was measured through the morphology of the nuclei and kinetoplasts stained with DAPI (Figure 1A). *T. brucei* TREU927 procyclic forms with 2N2K configuration were used to estimate the duration of C phase using Equation 1 (19), estimated as 0.82 h or 0.096 cell cycle unit (ccu). We found 6.99 ± 1.13% 2N2K procyclics from an assay carried out in biological triplicate (Figure 1A). To estimate the duration of the nuclear G2+M phases, *T. brucei* TREU927 procyclics were collected every 15 min in the presence of EdU until parasites containing two EdU-labeled nuclei (2N2K) were observed (Figure 1B). This pattern was first detected after 2 h, indicating that cells at the end of S phase required 2 h to proceed through G2 and M phases (Figure 1B). This assay was carried out in triplicate, and in all replicates, we found a parasite containing two EdU-labeled nuclei at the same time indicated. EdU-labeled parasites 1 h after an EdU pulse indicated the proportion of parasites able to replicate DNA (39 ± 2.7% of procyclics from *T. brucei* TREU927) (Figure 1C). Using this proportion and the estimated duration of the G2 + M + C phases, we were able to calculate the duration of S phase using Equation 2 (20). S phase was estimated to be 2.31 h or 0.272 ccu (Figure 1D). Finally, the duration of G1 phase was calculated as the difference between the sum of the durations of the other phases (S + G2 + M + C) and the doubling time. Thus, G1 phase duration was estimated to be 3.37 h or 0.397 ccu (Figure 1D).

These new precise values for the duration of G1 and S phase in *T. brucei* TREU927 share close similarities to those obtained for *T. brucei* Lister strain 427, which were also estimated by EdU incorporation (15) or by counter-flow elutriation (27). This similarity suggests that different strains of the same species (*T. brucei*) probably do not have significant differences in cell cycle lengths.

### The constitutive origins mapped in *T. brucei* are not sufficient to accomplish complete DNA replication within the S phase duratio

Using the newly estimated S phase duration, we estimated the minimum number of origins (*mo*) on each megabase chromosome of *T. brucei* through the application of Equation 3 (see materials and methods). During this estimation, we used two different values for the replication rate (*v* = 3.7 kb.min^-1^ and *v* = 1.84 kb.min^-1^) reported by different studies (3, 17), which resulted in two different values for *mo* (Table 1). These two values for *v* were calculated using a DNA combing technique, but each had a different approach (3, 17). Interestingly, another study developed in *T. brucei* mapped the position of activated origins by an MFA-seq technique, which matched ORC binding sites identified by ChIP-seq (16). This colocalization strengthens the assertion that these activated origins are indeed true origins and not possible products of other processes that generate DNA synthesis, such as DNA repair. However, the same study identified many more ORC binding sites than activated origins (16), which can be easily explained by the fact that MFA-seq (genome-wide analysis) is a technique that provides a result based on population analysis. Thus, the set of activated origins that matches with ORC binding sites can be classified as constitutive, and the remaining ORC binding sites are probably potential sites for the firing of non-constitutive origins (flexible or dormant) (6, 14).

**Table 1.** Comparison between the minimum numbers of origins (estimated using two different *v* values) and the constitutive origins detected by MFA-seq technique.

| *T. brucei* (TREU927) |  |  |  |  |
| --- | --- | --- | --- | --- |
| Chromosome | estimated chromosome size | minimum number of origins estimated using $\nu = 3.7$ kb.min<sup>-1</sup> | minimum number of origins estimated using $\nu = 1.84$ kb.min<sup>-1</sup> | constitutive origins mapped by MFA-seq |
| *I* | 1064,672 kb | 2 | 3 | 2 |
| *II* | 1193,948 kb | 2 | 3 | 3 |
| *III* | 1653,225 kb | 2 | 4 | 3 |
| *IV* | 1590,432 kb | 2 | 4 | 3 |
| *V* | 1802,303 kb | 2 | 4 | 3 |
| *VI* | 1618,915 kb | 2 | 4 | 2 |
| *VII* | 2205,233 kb | 3 | 5 | 2 |
| *VIII* | 2481,19 kb | 3 | 5 | 6 |
| *IX* | 3542,885 kb | 4 | 7 | 4 |
| *X* | 4144,375 kb | 5 | 9 | 6 |
| *XI* | 5223,313 kb | 6 | 11 | 8 |

S phase = 138.6 min (current study); replication speed = 3.7 kb.min<sup>-1</sup> (17), replication speed = 1.84 kb.min<sup>-1</sup> (3), number of constitutive origins per chromosome (16).

Comparing the two *mo* numbers estimated for each chromosome using Equation 3 with the number of constitutive origins, we easily notice that the *mo* estimated with *v* = 3.7 kb.min^-1^ shares close similarities with the number of constitutive origins (Table 1 and Figure 2A). However, the *mo* values estimated with *v* = 1.84 kb.min^-1^ are slightly above those represented by constitutive origins (Table 1 and Figure 2A), which makes us wonder which of the two *v* values would allow the constitutive origins complete replication within the S phase duration. Interesting, in the same study in which *v* = 1.84 kb.min^-1^ was estimated, the authors suggested an average inter-origin distance (IOD) of 148.8 kb, *i.e*., one origin per ∼ 148.8 kb. If we extrapolate this to all eleven chromosomes, we find an average number of origins per chromosome that is 3.1 times higher than the *mo* estimated with *v* = 1.84 kb.min^-1^, 4.5 times higher than the constitutive origins estimated by MFA-seq, and 5.9 times higher than the *mo* estimated with *v* = 3.7 kb.min^-1^ (Figure 2B). Although these data suggest that *T. brucei* uses a very large number of total origins to complete DNA replication, the average IOD estimated by DNA combing has a bias that impairs its reliability: how do we know if the molecules used for the calculation represent the entire genome? In other words, since the quantification of molecules is carried out in a population, it is possible that several origins from the same chromosomal fragment are measured, which would introduce a bias into the analysis. In this way, we decided at first not to use the average IOD as a priori information in our simulations, even though it corroborated our results. Of note, *T. brucei* spp. naturally show many more than 11 chromosomes (28), but only the sequences of the megabase-sized chromosomes are available in the TriTrypDB database (16). Thus, all subsequent analyses performed on *T. brucei* spp. will take into account only these chromosomes.

**Figure 2.**
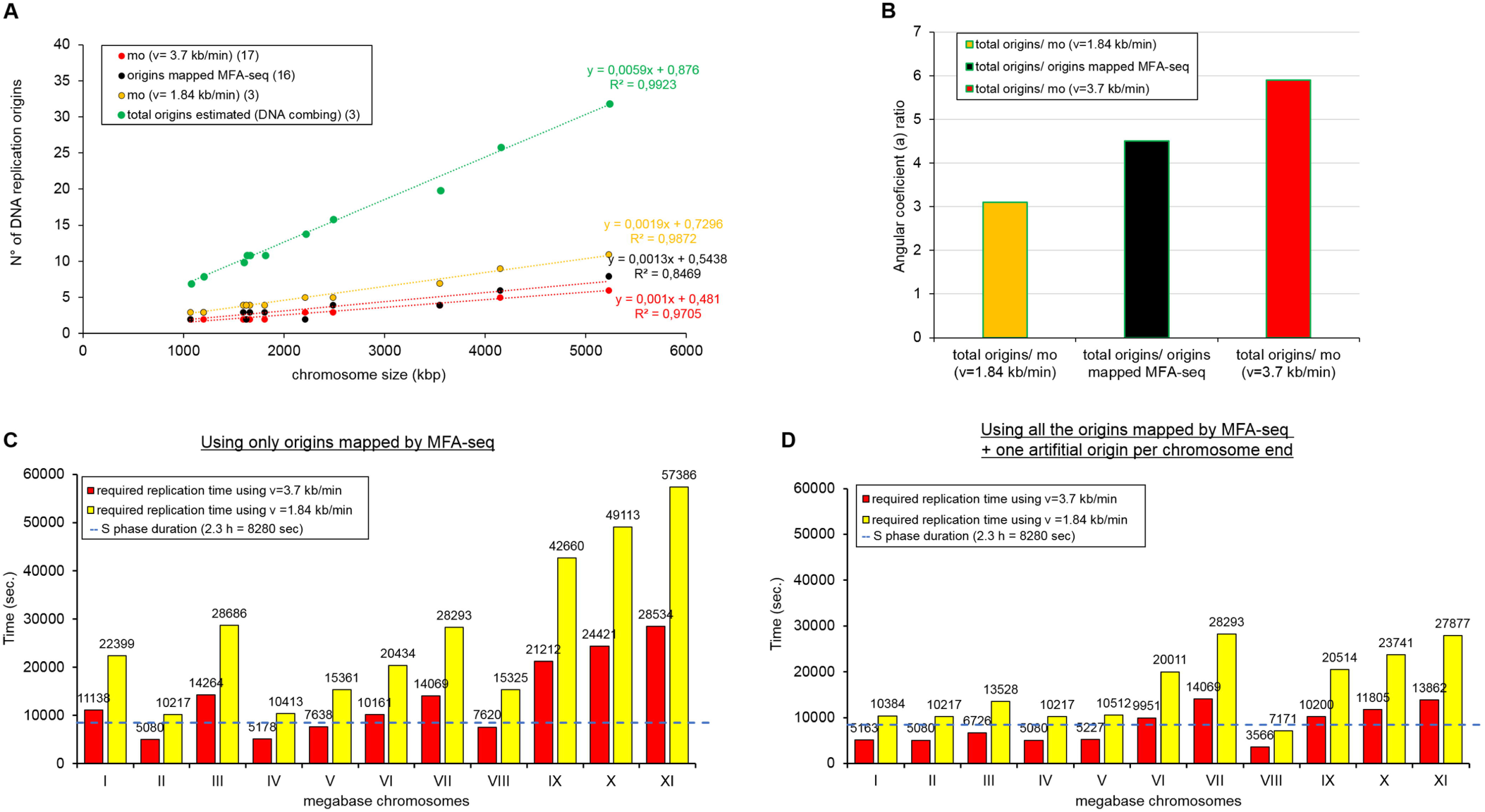
The constitutive origins mapped in *T. brucei* are not sufficient to accomplish full DNA replication within S phase. **A.** Graph showing positive correlations between chromosome length and the number of replication origins estimated by DNA combing (total origins) (green dots), number of origins estimated by MFA-seq (black dots), minimum origins (*mo*) estimated using *v* = 3.7 kb.min^-1^ (red dots), and *mo* estimated using *v* = 1.84 kb.min^-1^ (yellow dots). The trend lines for all groups, as well as the equations, are shown. **B**. Angular coefficient ratios between total origins and *mo* using *v* = 1.84 kb.min^-1^ (yellow bar = 3.1), total origins and origins estimated by MFA-seq (black bar = 4.5), and total origins and *mo* using *v* = 3.7 kb.min^-1^ (red bar = 5.9). **C.** Minimum time required for each *T. brucei* chromosome to complete DNA replication according to the positions of the origins mapped by MFA-seq (16), using two different values for the replication rate: *v* = 3.7 kb.min^-1^ (red) or *v* = 1.84 kb.min^-1^ (yellow). The dashed line represents the estimated S phase duration reported in this study (see Figure 1). **D**. To solve the bias generated by the possible presence of origins hidden in subtelomeric/telomeric regions, we repeated the assay shown in C by adding an artificial origin per chromosome end. Of note, each origin was localized 50 kb upstream of the chromosome end.

To determine if the given set of constitutive origins from *T. brucei* TREU927 (16) can complete the replication of each chromosome within the S phase duration, we developed another formula (Equation 4). This equation is able to calculate the minimum required time for DNA replication using only the set of constitutive origins under two conditions: *v* = 3.7 kb.min^-1^ or *v* = 1.84 kb.min^-1^ (see Materials and methods). When we used the fastest of these two estimations for *v* (3.7 kb.min^-1^), DNA replication was not completed within S phase duration since only four out of eleven chromosomes could accomplish DNA replication in this time frame (Figure 2C, Supplementary Table S1). These data may be contradictory since MFA-seq identified more origins than *mo* (Figure 2A); however, the results are easily justified due to the positions of the mapped origins (16), which do not allow an ideal configuration to optimize replication. Using *v* = 1.84 kb.min^-1^, the situation was even worse, as none of the eleven chromosomes allowed DNA replication within the S phase duration (Figure 2C).

It is worth mentioning that our *in silico* analyses used the value for full chromosome lengths available at TriTrypDB (www.tritrypdb.org). However, the lengths of *in silico* chromosomes may not reflect real *in vivo* chromosomal lengths in trypanosomatids since the annotation of some regions may not have been carried out with the necessary accuracy and details, such as for contigs containing subtelomeric and telomeric repeats (29, 30). Indeed, poorly annotated regions may affect also MFA-seq analysis since the localization of possible origins at subtelomeric/telomeric regions would be impaired. However, this would only be true for origins fired in late S phase, as those fired in early/middle S phase would potentially be detected because the upstream replisomes would have sufficient time to reach the innermost regions of the chromosomes. A genome-wide analysis of chromosome VI from *T. brucei* provides evidence for this argument (16). Even so, to address this critical point, we again carried out the simulations shown in Figure 2C but introduced one artificial origin in each chromosome end (each origin was localized 50 kb upstream of the chromosome end). We proposed to determine if the set of constitutive origins mapped by MFA-seq plus the artificial subtelomeric/telomeric origins could complete replication within the S phase duration. Surprisingly, it was not possible to complete DNA replication within the duration of S phase in the presence of artificial origins using a replication rate of *v* = 3.7 kb.min^-1^ or *v* = 1.84 kb.min^-1^ (Figure 2D, Supplementary Table S1). Using *v* = 3.7 kb.min^-1^, only five out of eleven chromosomes could accomplish DNA replication in the appropriate time. Using *v* = 1.84 kb.min^-1^, none of the eleven chromosomes could complete replication within the S phase duration.

### Dynamic model suggests an increase in replication initiation due to replication-transcription conflicts during S phase in *T. brucei*

Considering that trypanosomatids organize the majority of their genes into large polycistronic gene clusters (11, 31), this raises an issue: could an event related to transcription and replication (*e.g*., replication-transcription conflicts) contribute to the firing of flexible and/or dormant origins, thus increasing the pool of used origins, as previously suggested in other cell types (6, 9, 32)? To investigate the possible impact of replication-transcription conflicts on origin firing, we designed a stochastic dynamic model that constrained the availability of resources by establishing a limit for the number of available replisomes during simulation (parameter F, see materials and methods), as previously done in a similar model (21). Of note, in human cells, this role is played by the CDC45 protein (33). This model has a probability of replication origin firing modulated by the probability landscape reported in Supplementary Figure S1 (blue lines). Initially, for different values of F, we evaluated the model dynamics without the presence of transcription; in these simulations, we observed that there was a decrease in the mean inter-origin distance (IOD) as we increased the size of F (yellow line in Figure 3, Supplementary Table S2), with mean IOD values ranging from 558.5 kb (F = 10) to 70.3 (F = 100). Of note, the mean IOD calculated using putative constitutive origins as presented by Tiengwe et al. is equal to 631.4 kb; this means that all sets of results provided by the dynamic model were below this upper bound for the mean IOD. Interestingly, for F = 45, the mean IOD was 147.7 kb, which was almost the same as the 148.8 kb suggested in the previous study in which *v* = 1.84 kb.min^-1^ was estimated (3).

**Figure 3.**
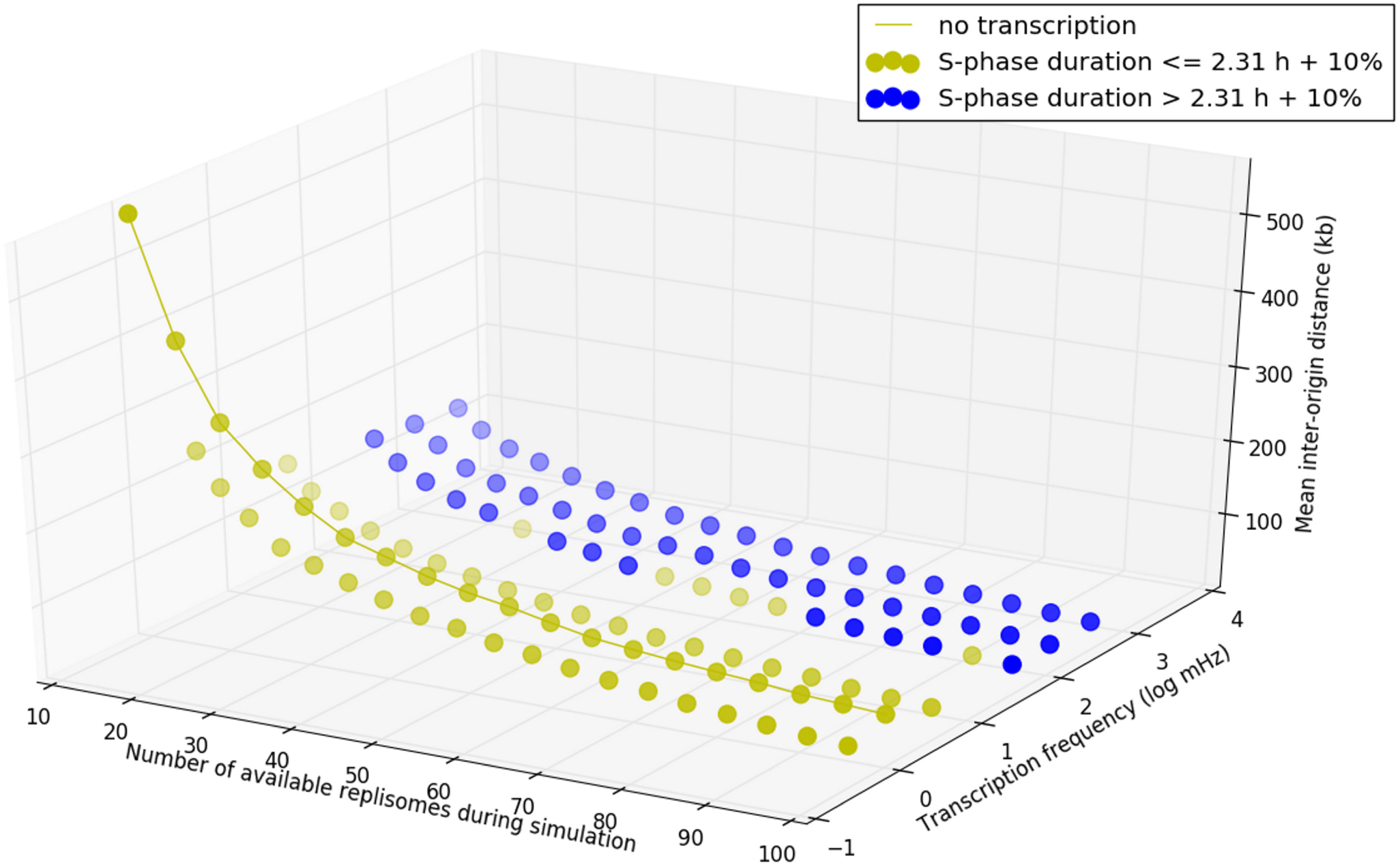
Computational simulations suggest that replication-transcription conflicts increase the activation of origins. **A.** Results of simulations with a stochastic dynamic model show the mean inter-origin distance (which is inversely proportional to the activation of origins) as a function of the transcription frequency and the number of available replisomes during a simulation (parameter F). The yellow line represents sets of simulations without the presence of constitutive transcription. For a given F value, yellow and blue dots represent sets of simulations that, on average, had DNA replication durations up to 10% and more than 10%, respectively, higher than those verified for simulations without transcription.

In sequence, we evaluated the impact of increasing frequencies of constitutive transcription on replication-transcription conflicts and on S phase duration. These simulations indicated a reduction in the mean IOD (which is inversely proportional to the number of replication initiations) in response to increasing values of the parameter F and to increasing values of the constitutive transcription frequency (Figure 3, Supplementary Table S2). Even simulations with low transcription frequency led to at least a 2-fold reduction in the mean IOD relative to with simulations without transcription (Figure 3, yellow line). Additionally, the system displayed a robustness of S phase duration for moderate amounts of transcription frequency (Figure 3A, yellow points).

Thus, under standard conditions, replication-transcription conflicts might be responsible for a decrease in the estimated mean IOD, making the parasite fire a pool of origins greater than the minimum required to complete S phase. To verify the reliability of this *in silico* assay, we decided to validate these predictions experimentally, starting by ascertaining the presence of transcription during S phase.

### Polycistronic transcription in *T. brucei* TREU927 is carried out throughout the cell cycl

To confirm the presence of transcription during S phase of *T. brucei* TREU927, the parasites were subjected to an *in vitro* transcription methodology (nuclear run-on assay) coupled with DNA content analysis using DAPI. This methodology enabled us to analyze the transcription of nascent RNAs transcribed mainly by RNAP II during different cell cycle phases. In this way, we analyzed transcriptional activity according to the DNA content profile in a wild type population of *T. brucei* (control) (Figure 4A - left). We observed transcription activity throughout the cell cycle (G0/G1, S, and G2/M), indicating that there is nascent RNA synthesis in all phases of the cell cycle of *T. brucei* TREU927 (Figure 4A, green group). To validate the data, we pretreated the parasites with α-amanitin, an irreversible transcription inhibitor (Figure 4A - right). The α-amanitin treatment caused a drastic decrease in the nascent RNA signal (Figure 4A, green group). This result confirms that RNAP II transcribes nascent polycistronic RNA during all the phases of the cell cycle, including S phase. Figure 4B shows the DNA content profiles of both analyzed groups: control and α-amanitin-treated.

**Figure 4.**
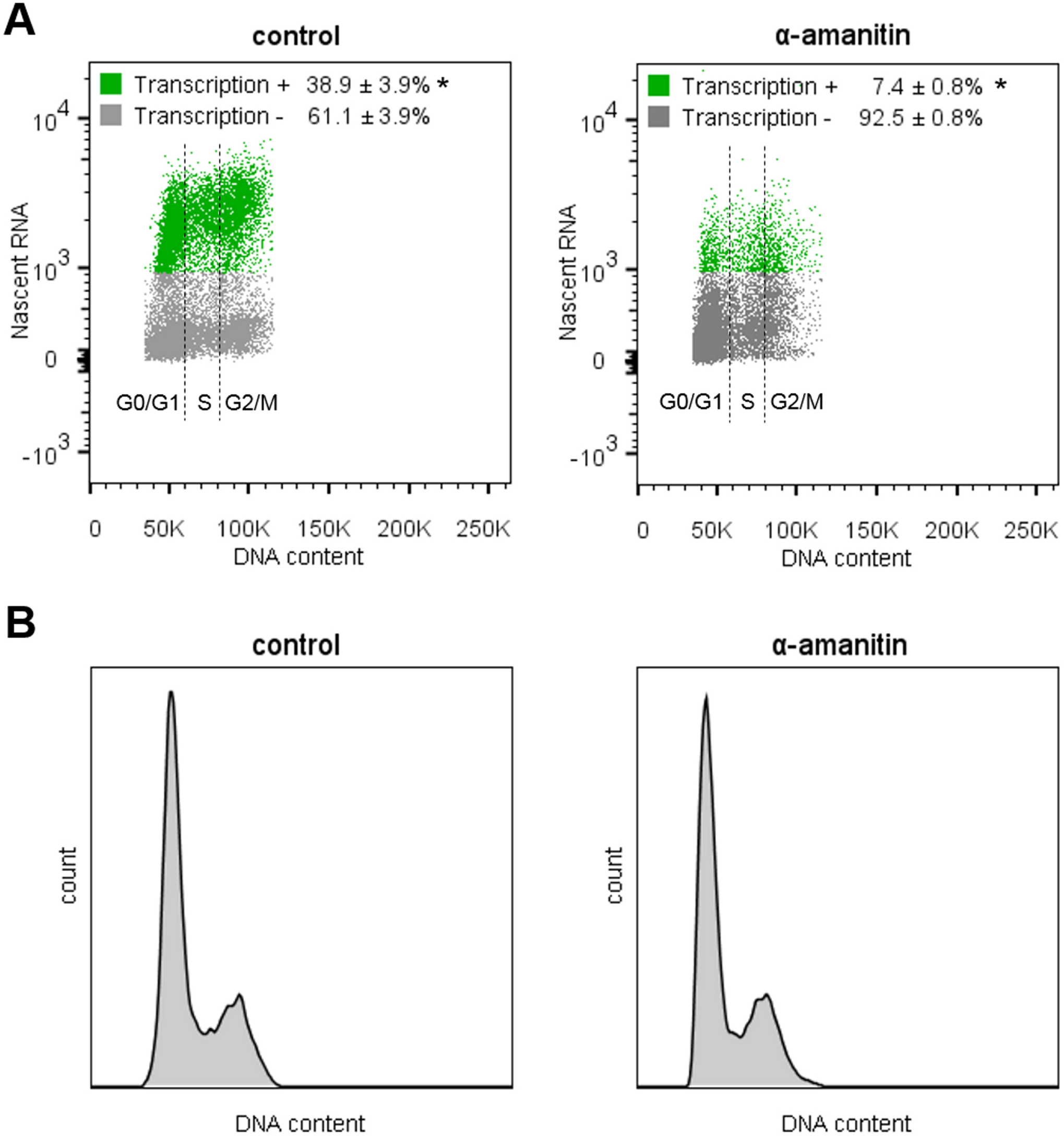
Transcription landscape throughout the *T. brucei* TREU927 cell cycle and simulations including transcription interference. **A.** Distribution of the population according to DNA content and RNA synthesis. Left – Wild-type (control) population showing transcriptional activity = 38.9 ±3.9% (green). Right – After transcription inhibition using α-amanitin, the parasites performing transcription decreased to 7.4 ±0.8% (green). The cell cycle phases (G0/G1, S, and G2/M) are indicated. Green – transcription-positive, gray – transcription-negative. **B.** Histograms showing the largely unchanged DNA content profiles for the control (left) and α-amanitin treated groups (right). These assays were carried out in biological triplicate, and 20,000 parasites (n = 20,000) were counted in each analysis (* represents p < 0.01 using Student’s t-test).

### Endogenous fluorescent foci of γH2A and R-loops are detected during S/G2 phase

Exponential *T. brucei* procyclic forms were used in indirect immunofluorescence assays (IIF) to measure the endogenous levels of DNA breaks and R-loops in different cell cycle phases; both are consequences of replication-transcription conflicts (8). Importantly, morphological distinction of the cell cycle phases was performed based on DAPI-stained nuclei/kinetoplasts, allowing us to differentiate cells with nuclei in G1/early S, late S/G2, mitosis, and cytokinesis in *T. brucei* (Figure 5) (27, 34). To analyze the endogenous levels of DNA breaks, we measured the fluorescence intensity of the histone γH2A, which is phosphorylated predominantly in the presence of DNA lesions and is commonly used as a DNA break biomarker (34–36). It is worth mentioning that in mammals, the phosphorylation resulting from DNA breaks happens in the variant histone H2AX, which generates γH2AX. Therefore, γH2AX is commonly used as a DNA break biomarker. However, the presence of this variant histone has not been reported in trypanosomatids or yeast (34, 37). Several studies indicate that the canonical phosphorylated histone (γH2A) plays the role of γH2AX in these organisms, *i.e*., increasing *in vivo* in response to DNA lesions (34, 38, 39).

**Figure 5.**
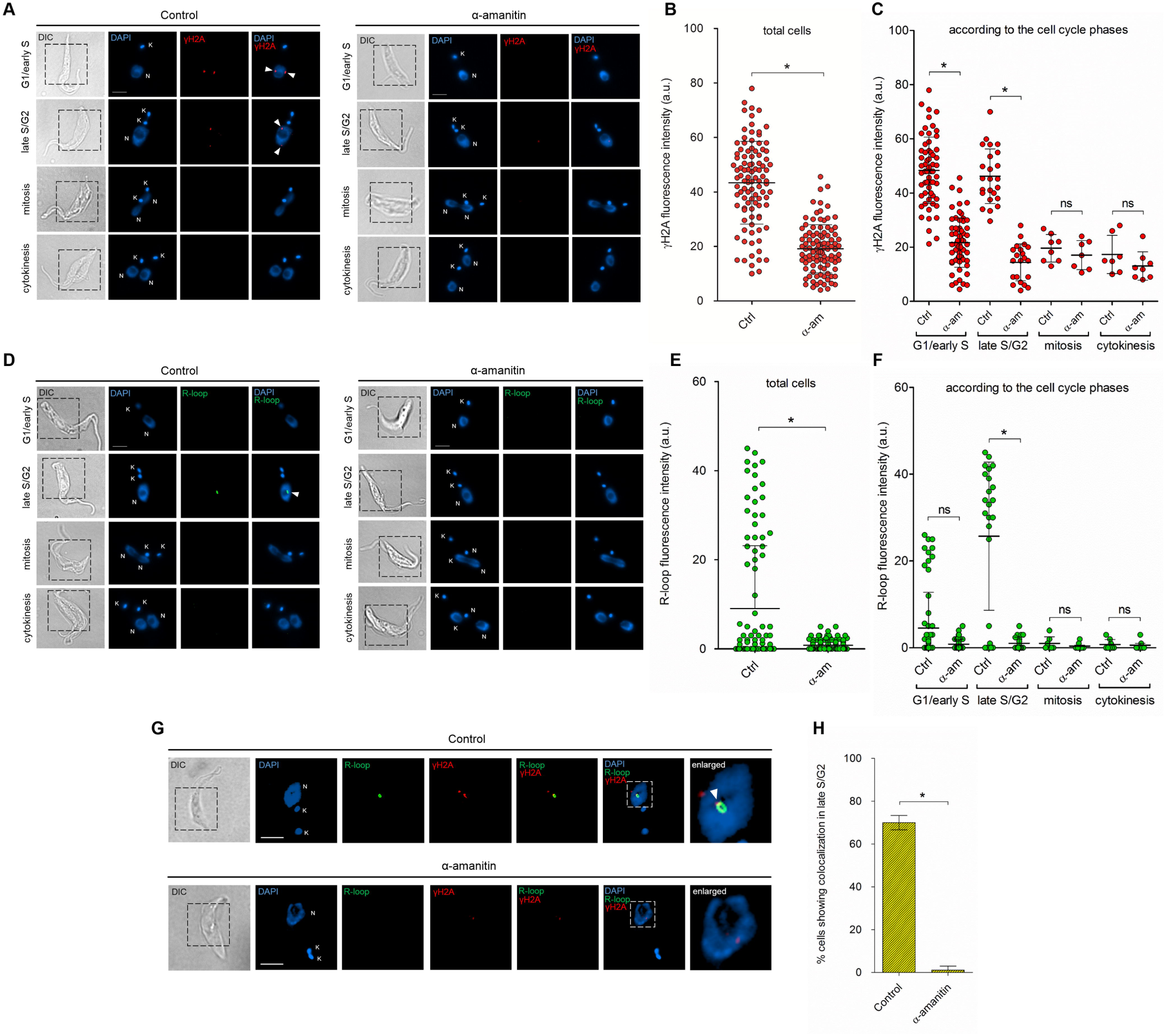
Distribution profile of γH2A and R-loops throughout the cell cycle in the presence and absence of transcription. **A**. (left) The endogenous γH2A foci shown by nontreated (control) parasites suggest the presence of DNA lesions, mainly during G1/early S and late S/G2. (right) These γH2A foci decreased after transcription inhibition (α-amanitin treated). **B.** Graph showing the γH2A fluorescence intensity (red) per cell in total cells and **C**. according to the cell cycle phase analyzed. Errors bars indicate SD. The difference observed was statistically significant using Student’s t-test (* = p < 0.001) for a biological triplicate assay (n = 100). **D**. (left) The control parasites showed endogenous R-loops foci, predominantly during late S/G2. (right) After transcription inhibition, these R-loop foci decreased. **E**. Graph showing the R-loop fluorescence intensity (green) per cell in total cells and **F**. according to the cell cycle phase analyzed. Errors bars indicate SD. The difference observed was statistically significant using Student’s t-test (* = p < 0.01) for a biological triplicate assay (n = 100). **G**. Representative confocal images showing colocalization (white triangle) between γH2A (red) and R-loop (green) during late S/G2 in the control group. After transcription inhibition, this colocalization virtually disappeared. **H**. The graph shows that 69.9 ± 3.35% of parasites (in late S/G2) showed colocalization, which fell to 1.1 ±1.9% after transcription inhibition. Errors bars indicate SD. The difference observed was statistically significant using Student’s t-test (* = p < 0.01) for a biological triplicate assay (n = 100 parasites).

*T. brucei* shows endogenous levels of DNA breaks represented by a few γH2A fluorescent foci (red), predominantly throughout the G1, S and G2 phases (Figure 5A). These foci became rare during the mitosis and cytokinesis phases (Figure 5A), which can be explained by the fact that, when DNA breaks occur during S phase, their repair takes place predominantly in the late S or G2 phases via the homologous recombination (HR) pathway (39, 40). Alternatively, HR repairs one-ended DSBs that arise following replication fork collapse throughout S phase (40). Both situations corroborate the γH2A fluorescent levels naturally observed in *T. brucei*, which was also shown in another study (34). In addition, when we inhibited transcription via treatment with α-amanitin, we observed a decrease in γH2A fluorescence basal levels during G1/early S and late S/G2. Significant differences in the fluorescence intensities of each individual parasites that were either not treated (control) or treated with α-amanitin were measured and are represented in the graphs (Figure 5B). Measurements according to the cell cycles phases are also shown in Figure 5C.

An important and pertinent observation that may be raised relates to the immediate phosphorylation of H2A and its dependence on transcription. In other words, how do we know if the immediate nondetection of γH2A fluorescence levels represents a bias due to the absence of transcription? To address this critical point, which is inherent to this assay, we induced DSBs using 50 Gy of ionizing radiation (IR) in the presence or absence of transcription (Supplementary Figure S8). We did not detect a decrease in γH2A fluorescence intensity after inducing DSBs in the absence of transcription. Thus, we can state that the immediate phosphorylation of H2A did not depend directly on nascent RNAs that were eventually inhibited during the period of treatment using α-amanitin (Supplementary Figure S8). Therefore, we can infer that transcription-dependent DNA breaks are generated endogenously in *T. brucei* TREU927, probably during S phase and remaining until G2, when the parasite repairs the damage and γH2A phosphorylation is removed (Figure 5A-C). Of note, this was not detected in parasites that were treated with 50 Gy because IR reaches the parasites in all cell cycle phases instantaneously, generating DSBs in all phases (Supplementary Figure S8).

To measure natural R-loop foci, we used a specific antibody (S9.6) that recognizes DNA:RNA hybrids (41). R-loops are triple-stranded nucleic acid structures composed of a DNA:RNA hybrid and a single-stranded DNA (ssDNA), and their generation is related to various factors, among them replication-transcription collisions (8, 42, 43). Similar to γH2A, *T. brucei* shows natural levels of R-loops represented by green fluorescent foci throughout the S and G2 phases, which become rare during mitosis and cytokinesis (Figure 5D and F). Interestingly, we have no evidence to speculate how often these R-loops are formed nor in which cell cycle phase these structures are resolved. However, as most of the mechanisms for the resolution of R-loops are described as acting throughout the cell cycle (44–46), it is intuitive to propose that R-loops are solved as soon as they are generated to avoid possible genome instability. Moreover, when transcription is inhibited by treatment with α-amanitin, we observed a decrease in R-loop fluorescent foci, which is indicated by the fluorescent images and graphs (Figure 5D-F). Thus, we may state that the generation of R-loops is transcription-dependent and is solved before cells enter mitosis. Additionally, to verify the specificity of the antibody used, we treated *T. brucei* parasites with RNase H (a ribonuclease that catalyzes the cleavage of RNA from a DNA:RNA substrate), and we detected a statistically significant decrease in R-loop fluorescent foci (Supplementary Figure S9).

### γH2A and R-loop foci colocalize in late S/G2 phase, and this colocalization disappears after transcription inhibition

Next, to determine if DNA damage is, in fact, a consequence of replication-transcription conflict, we asked whether γH2A and R-loops colocalize since both are observed in the same cell cycles phases (from S to G2) and are virtually transcription-dependent. For this, we performed IIF colocalization assays using confocal microscopy. We observed colocalization (indicated by the white triangle) only in late S/G2 cells, represented by cells with a 1N2K configuration (Figure 5G). This indicates that γH2A is preferentially accumulated at R-loop-enriched foci (Figure 5G). The percentage of cells in late S/G2 showing this colocalization was 69.9 ±3.35%. When we inhibited transcription via treatment with α-amanitin, colocalization decreased, and the percentage fell to 1.1 ±1.9% (Figure 5H). This result suggests that γH2A and R-loops foci, besides being transcription-dependent, are generated from the same location. We speculate that this colocalization occurs probably due to replication-transcription conflicts.

### *T. brucei* activates fewer replication origins in the absence of transcription

To experimentally compare the percentage of activated DNA replication origins in the presence (control) and in the absence of transcription (α-amanitin), as well as to validate the predictions of our stochastic computational models, we used a DNA combing approach. This technique allows the visualization of replication origins in replicated DNA molecules stretched on coverslips via the subsequent incorporation (short-pulse) of halogenated thymidine analogs, namely, IdU (red) and CldU (green). The DNA molecules are also labeled with anti-DNA (blue). Figure 6A shows the patterns we looked for during our analyses. Figure 6B shows some of the patterns (chromosome fibers) found in our analyses. From a group of 60 DNA molecules (n=60), we were able to detect origin and termination regions, as well as the direction of the replication fork. In the control group, we measured 56.6 ±9.4% origins, while this value decreased to 26.6 ±4.7% in the α-amanitin-treated group (Figure 6C).

**Figure 6.**
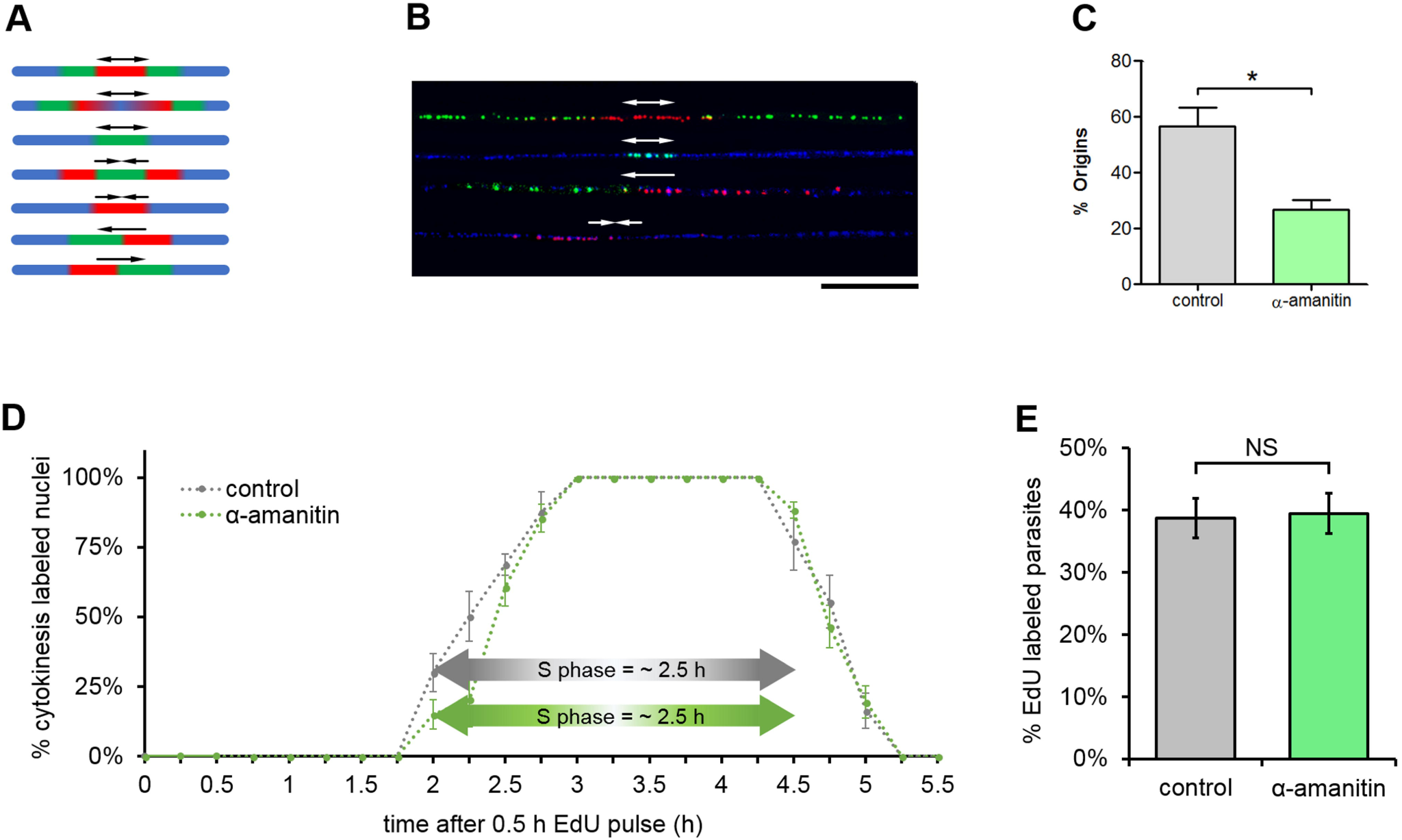
After transcription inhibition, *T. brucei* activates fewer replication origins to maintain the robustness of S phase. **A.** Patterns that we looked for during our DNA combing analysis. The arrows represent the fork direction. From top to bottom, the patterns represent origin, origin, origin, termination, termination, replication fork, and replication fork. **B.** Some representative chromosome fibers found in our analysis. The arrows represent the fork direction. Bar = 20 μm. We detected origin and termination regions, as well as activated replisomes in both directions. **C.** Graph showing that the percentage of origins decreased from 56.65 ±9.4% (control) to 26.67 ±4.7% (α-amanitin treated). The percentage of origins was calculated for a group of DNA molecules that incorporated both thymidine analogues (CldU and IdU); (*) p < 0.06 using Student’s t-test, n = 60 chromosome fibers. **D.** Estimation of S phase length by measurement of cytokinesis-labeled nuclei. After performing a 15-min EdU pulse, we quantified the percentage of cytokinesis-labeled nuclei every 15 min in the control and α-amanitin-treated groups; the results indicated nonsignificant differences in S phase length after transcription inhibition. The bars represent the SD from an assay carried out in biological triplicate (n = 20 cells for each time point analyzed, totaling n = 460 cells). **E**. The percentage of parasites able to uptake EdU after α- amanitin treatment remained largely the same (38.7 ±3.2% for control and 39.5 ±3.2% for the α- amanitin-treated group). This demonstrates that transcription inhibition did not impair DNA replication in the measured period. The difference observed was not statistically significant using Student’s t-test (NS = not-significant) for a biological triplicate assay (n = 569 cells).

Corroborating our stochastic computational simulations carried out previously (comparing these results with those in Figure 3 and Supplementary Table S2), this assay suggests that fewer origins are activated in the absence of transcription, which points to the contribution of the transcription machinery in the firing of origins, possibly as a result of replication-transcription conflicts.

### S phase duration remains the same even in the absence of transcription

To quickly measure S phase length during transcription inhibition, avoiding possible transcription-dependent interference, we decided to use an approach that does not rely on the measurement of other phases of the cell cycle, such as those performed in Figure 1. To this end, we followed an established assay in trypanosomatids (15), which directly yields kinetic determinations of S phase through the measurement of cytokinesis (2N2K)-labeled nuclei. In this assay, we performed a 15-min EdU pulse to subsequently measure the cytokinesis of EdU-labeled nuclei until the EdU signal disappeared. There was an initial period in which cytokinesis was not labeled, and this period finished with the onset of cytokinesis-labeled nuclei that sharply increased up to 100% and stayed at this level for a defined period of time, yielding estimates of S phase length: ∼ 2.5 h for both the control and α-amanitin-treated groups (Figure 6D). Both cytokinesis labeling curves were practically coincident, which means that S phase duration was not affected by transcription inhibition (Figure 6D). These assays were performed in biological triplicate (n = 20 for each time point), and the error bars represent the SD. Of note, during these kinetic experiments, the parasite populations grew exponentially and in a steady-state manner. Despite the exponential growth of the populations, labeling indexes displayed approximate linear increases since the measures were carried out in a period shorter than the doubling time for *T. brucei* TREU927. In addition, the capacity of the parasites for EdU uptake after α-amanitin treatment remained largely the same (Figure 6E). This demonstrates that short-term transcription inhibition does not affect DNA replication, in agreement with the dynamic model prediction of robustness in S phase duration with respect to the absence/presence of transcription (Figure 3, Supplementary Table S2).

## DISCUSSION

The precise duration of S phase in *T. brucei* TREU927 estimated using EdU (Figure 1) allowed us to estimate the minimum number of origins (*mo*) required to replicate an entire chromosome during S phase (see Materials and methods). After comparing the *mo* values estimated using the two different replication rates available in the literature [*v* = 3.7 kb.min^-1^ (17); *v* = 1.84 kb.min^-1^ (3)] with the constitutive origins mapped by MFA-seq, we asked if these origins (the constitutive ones) were sufficient to accomplish DNA replication within the S phase duration (Figure 2A, B). Through stochastic computational models, we determined that these origins were not sufficient to accomplish complete DNA replication within the S phase duration, even in the presence of artificial subtelomeric/telomeric origins (Figure 2C, D). This demonstrates that, regardless of the hypothetical presence of replication origins within subtelomeric/telomeric regions, and even if the number of constitutive origins mapped by MFA-seq per chromosome is greater than the *mo* estimated with *v* = 3.7 kb.min^-1^, the positioning of origins does not permit the replication of all chromosomes within the S phase duration. A parsimonious hypothesis that could satisfactorily explain this peculiarity is the activation of non-constitutive origins. These origins may be flexible or dormant, although the presence of both is not mutually exclusive, *i.e*., the presence of replication stress that activates dormant origins does not exclude the possibility that the flexible origins are also activated in a stochastic manner. One piece of evidence supporting this hypothesis is that the MFA-seq data found many more ORC binding sites than replication peaks, suggesting the presence of non-constitutive origins in *T. brucei* TREU927 (16). In addition, a study that used a DNA combing approach, as previously mentioned (17), also suggested the presence of flexible origins in this parasite.

Based on these features and considering that the precise S phase duration estimated previously (Figure 1) is robust, as proposed for other cell types (21, 47), simulations with our stochastic computational models suggest that the presence of replication-transcription conflicts might lead to an increase in origin firing, which in turn helps to maintain the robustness of the S phase duration (Figure 3, Supplementary Table S2). Analyzed together, these simulations suggest the presence of a mechanism responsible for a decrease in the mean *IOD* estimated under standard conditions, making the parasites fire a pool of origins greater than the minimum required to complete S phase. As trypanosomatids organize the majority of their genes into large polycistronic gene clusters (31), we hypothesized that some event related to transcription, which would generate replication stress, could contribute significantly to the activation of non-constitutive origins. Our computational models demonstrated that this hypothesis is fully possible since transcription during S phase directly impacts the number of origins required to complete replication in the estimated time frame (Figure 3, Supplementary Table S2). As transcription occurs during S phase (Figure 4), both replisomes and RNAP may operate concomitantly in the same region, making it inevitable that these two processes interfere with each other, generating collisions. The consequences of these collisions represent a high risk for most cells, as they may give rise to replication fork stalling or collapse, which is often accompanied by DNA damage, recombination or, in some cases, cell death (8, 48–51).

We provide evidence here that the endogenous levels of DNA damage shown by *T. brucei* can be a consequence of transcription activity (Figure 5). These levels of DNA damage were indirectly shown by the presence of fluorescent foci of the histone γH2A, predominantly during G1/early S and late S/G2 and decreasing after transcription inhibition (Figure 5A-C). The γH2A foci observed are in agreement with other studies in trypanosomatids (34, 52) and in cancer cells, which showed endogenous levels of γH2AX at transcription start sites (53, 54). However, not all γH2A foci decreased after transcription inhibition (Figure 5A-C), which makes it clear that transcription activity (possibly due to conflicts with replication) contributes to the generation of endogenous DNA lesions, but it is not its exclusive cause. Alternatively, the extension of stalled forks over longer time periods and the erosion of uncapped telomeres may generate one-ended DSBs, which could also give rise to the formation of endogenous γH2A(X) foci (40, 55, 56). In general, one-ended DSBs are repaired during S phase by a mechanism similar to homologous recombination (HR) called break-induced replication (BIR) (56), while common two-ended DSBs generated by replication stress are repaired by HR, but in late S and G2 phases (39, 40, 57, 58). This may explain the low γH2A signal during mitosis and cytokinesis in our results (Figure 5A, C and Supplementary Figure S8), *i.e*., it is likely that *T. brucei* efficiently repairs its endogenous DNA damage during S/G2. Such a proposal has previously been supported in this parasite after the induction of DSBs by ionizing radiation (IR), where repair was carried out quickly in late S/G2 within approximately 6 h (39). Thus, we can infer that endogenous DNA damage does not appear to be a great obstacle for this parasite to continue to proliferate, given the efficient DNA repair that most trypanosomatids possess (39, 52, 59, 60).

Furthermore, we detected fluorescent R-loop foci predominantly in late S/G2 (Figure 5D- F and Supplementary Figure S9). Curiously, these R-loop foci were found only near the nucleolus, which corresponds to the area that is less heavily stained by DAPI within the nucleus (61). This can be easily explained by the high levels of transcription carried out by RNAP II, which, in *T. cruzi*, largely takes place in a similar position relative to the R-loops we observed (61). Obviously, this does not imply that R-loops are present only in this region, but since transcription is more abundant there, fluorescent foci become more pronounced, while the other possible R-loop sites probably are hidden due to the low sensitivity of this technique for their detection. Treatment with exogenous RNase H, an enzyme that solves these structures (45, 62), confirmed that the foci we observed were indeed R-loops (Supplementary Figure S9). Moreover, after transcription inhibition, as expected, these R-loops foci completely disappeared (Figure 5D-F). In model eukaryotes, some studies have suggested that R-loops occur at a low frequency during replication (63, 64), and the disturbance of replication or transcription favors their creation or stabilization, which leads to increased DNA damage (63–65). Interestingly, we detected partial colocalization between DNA lesions and R-loops in 70 ± 3.35% of the parasites in late S/G2 (Figure 5G-H), which corroborates the results of some studies carried out in other cell types (66–68). This result allows us to establish a correlation between DNA damage and R-loops in trypanosomatids but does not allow us to establish a causal relationship. In other words, further studies are required to answer whether DNA lesions are generated in response to the presence of R-loops, or whether lesions are generated concomitantly with them.

An interesting point is that R-loops should also be mediated in an origin-independent replication mechanism, as demonstrated in yeast (66), human mitochondria (67), and prokaryotes (68, 69). If this is the case, R-loops could prime DNA replication in loci where RNAP activity is impaired, such as DNA repetitive sequences (70). Further studies are necessary to investigate whether this hypothesis is true in trypanosomatids. Regardless, our results highlight transcription as an important player in the modulation of endogenous R-loops, as previously suggested in bacteria (42, 71), yeast (72), *C. elegans* (72), and human cells (72, 73).

Using a DNA combing approach, we demonstrated that the percentage of activated origins decreases after transcription inhibition, probably due to the absence of origin firing (Figure 6C). This result suggests that transcription activity allows the firing of non-constitutive origins. However, our assays could not distinguish between passive (stochastic) or active (triggered by DNA replication impairment) origin firing. Thus, the mechanism by which non-constitutive origins are activated is an open question that requires further experiments. We also demonstrated that after transcription inhibition, S phase duration remained the same, at least after a short period (Figure 6D). This result confirms the robustness of S phase duration, as previously suggested by other studies (47, 74). Several studies have demonstrated the activation of dormant origins following replication stress to ensure the completion of DNA replication near stalled or collapsed replication (6, 14, 75–77). However, none of these studies indicated that replication-transcription conflicts contributed to the firing of origins, which, in model eukaryotes, is an apparently well-regulated event (6). Based on this, we have proposed that DNA damage is generated after replication and transcription find each other in conflicting directions, which signals to initiate the phosphorylation of a nearby histone H2A, either by ATR or ATM kinases (14, 34, 78). Moreover, the hybrid R-loop structure is likely formed and maintained in the collision region. Concomitantly, a nearby origin is fired (in an active or passive manner) to complete replication and maintain the robustness of S phase (Figure 7).

**Figure 7.**
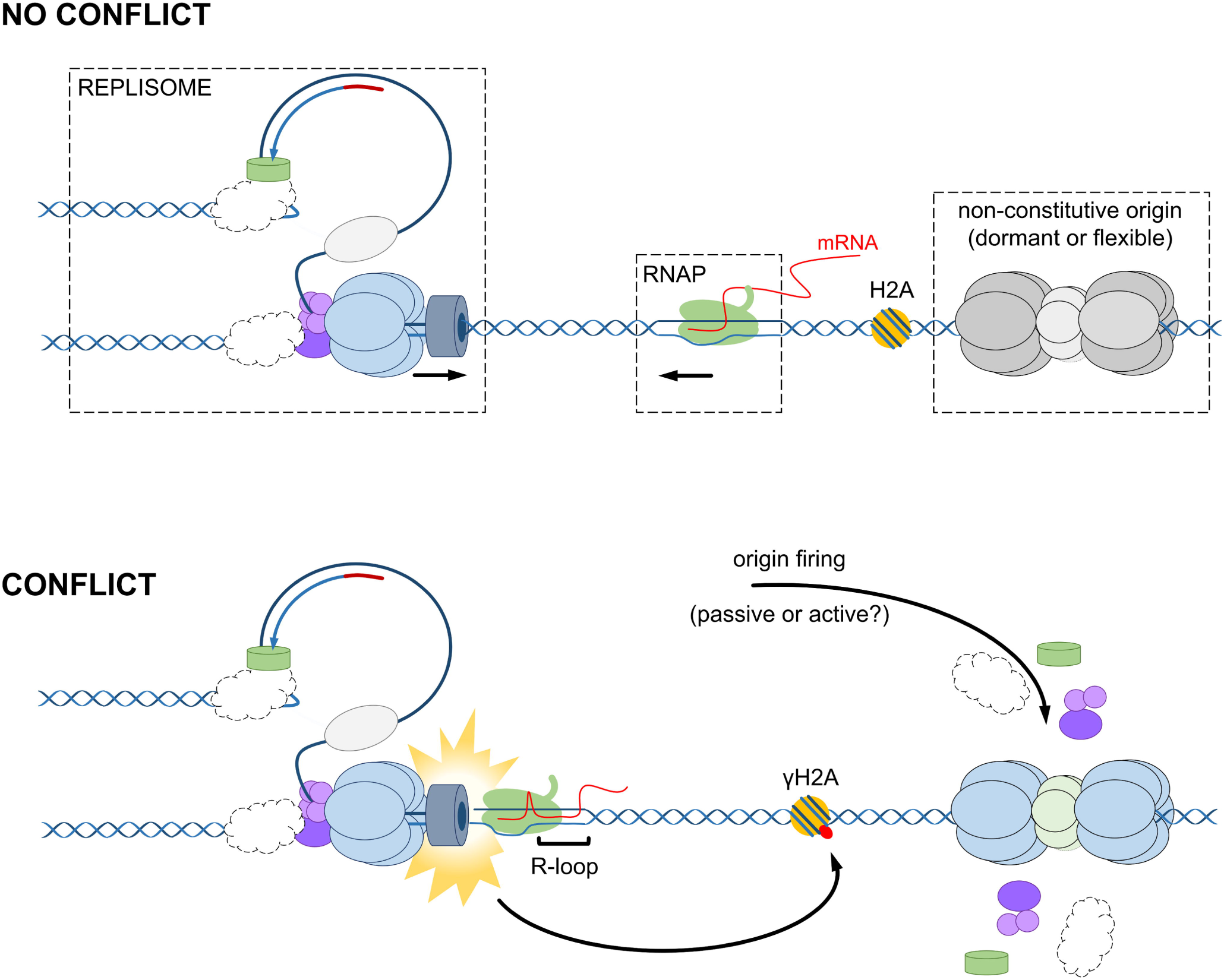
Schematic elucidating the possible consequences of replication-transcription conflicts. Genomic instability, represented in this case by R-loops and γH2A, is probably one of the worst consequences of these conflicts. To mitigate this problem, non-constitutive origins are activated, which help to maintain the robustness of S phase, contributing to the maintenance of DNA replication. Of note, the activation of non-constitutive origins may occur in an active (triggered by DNA replication impairment) or passive (stochastic) manner.

It is worth mentioning that, as a group, trypanosomatids emerged late on the evolutionary scale and are one of the most successful groups of specialist parasites on earth (14, 79, 80). This success probably occurred because these organisms proliferate rapidly, which increases the onset of mutation frequency due to multiple DNA replication events in a relatively short period (81–83). Whether replication and transcription in conflicting directions lead to increased mutagenesis, promoting the faster adaptive evolution of core genes, as demonstrated in *Bacillus subtilis* (84), is a question for debate that requires further studies. In conclusion, our results suggest that the conflict between transcription machinery action and replication is the main contributor to increased origin usage. The use of this entire pool of origins is necessary to maintain the robustness of S phase duration and seems to be of paramount importance since it also contributes to the maintenance of DNA replication, allowing the survival of this divergent parasite.

## ACKNOWLEDGMENTS

The authors thank the Fiocruz Network of Technology Platforms for use of the flow cytometry facility (RPT08L) at Carlos Chagas Institute – Fiocruz/PR. In addition, the authors are grateful to Dr. Richard McCulloch at the Wellcome Centre for Molecular Parasitology, University of Glasgow, for the antisera (anti-γH2A) he provided and for important observations and discussion. Moreover, we are grateful to the São Paulo Research Foundation (FAPESP) and Center of Toxins, Immune Response and Cell Signaling (CeTICS) for their support. The authors thank the Program for Technological Development in Tools for Health-RPT-FIOCRUZ for the use of the flow cytometry facility (RPT08L) at Carlos Chagas Institute - Fiocruz/PR.

